# Gene expression polymorphism underpins evasion of host immunity in an asexual lineage of the Irish potato famine pathogen

**DOI:** 10.1101/116012

**Authors:** Marina Pais, Kentaro Yoshida, Artemis Giannakopoulou, Mathieu A. Pel, Liliana M. Cano, Ricardo F. Oliva, Kamil Witek, Hannele Lindqvist-Kreuze, Vivianne G. A. A. Vleeshouwers, Sophien Kamoun

**Affiliations:** The Sainsbury Laboratory, Norwich Research Park, Norwich, United Kingdom; Graduate School of Agricultural Science, Kobe University, Japan; Wageningen UR Plant Breeding, Wageningen, The Netherlands; Institute of Food and Agricultural Sciences, Department of Plant Pathology, Indian River Research and Education Center, University of Florida, Fort Pierce, FL, USA; Genetics and Biotechnology Division, International Rice Research Institute, Los Banos, Philippines; International Potato Center, Lima, Peru

## Abstract

Outbreaks caused by asexual lineages of fungal and oomycete pathogens are an expanding threat to crops, wild animals and natural ecosystems (Fisher et al. 2012,Kupferschmidt 2012). However, the mechanisms underlying genome evolution and phenotypic plasticity in asexual eukaryotic microbes remain poorly understood (Seidl and Thomma 2014). Ever since the 19^th^ century Irish famine, the oomycete *Phytophthora infestans* has caused recurrent outbreaks on potato and tomato crops that have been primarily caused by the successive rise and migration of pandemic asexual lineages (Cooke et al. 2012, Yoshida et al. 2013,Yoshida et al. 2014). Here, we reveal patterns of genomic and gene expression variation within a *P. infestans* asexual lineage by compared sibling strains belonging to the South American EC-1 clone that has dominated Andean populations since the 1990s (Forbes et al. 1997, Oyarzun et al. 1998, Delgado et al. 2013, Yoshida et al. 2013, Yoshida et al. 2014). We detected numerous examples of structural variation, nucleotide polymorphisms and gene conversion within the EC-1 clone. Remarkably, 17 genes are not expressed in one of the two EC-1 isolates despite apparent absence of sequence polymorphisms. Among these, silencing of an effector gene was associated with evasion of disease resistance conferred by a potato immune receptor. These results highlight the exceptional genetic and phenotypic plasticity that underpins host adaptation in a pandemic clonal lineage of a eukaryotic plant pathogen.

---

Sexual reproduction is an important source of genetic diversity yet many asexual fungal and oomycete pathogens can rapidly adapt to their host environment and cause destructive pandemics (de Jonge et al. 2013, Farrer et al. 2013, Castagnone-Sereno and Danchin 2014,Seidl and Thomma 2014). Filamentous plant pathogens, including the soil-borne wilt fungus *Verticillium dahliae,* the rice and wheat blast fungus *Magnaporthe oryzae,* several rust fungi and *Phytophthora* spp. have widespread clonal populations that are a threat to managed and natural ecosystems as well as global food security (Leonard and Szabo 2005, Ebbole 2007, Gruünwald et al. 2008, Goellner et al. 2010, Cooke et al. 2012, de Jonge et al. 2013, Hubbard, Badran et al. 2015, Short et al. 2015,Islam et al. 2016). The establishment of these asexual populations is favored by agricultural practices such as monoculture and year-round crop production (Stukenbrock and McDonald 2008). Therefore, asexual reproduction in plant pathogens is an important feature with serious epidemiological and agronomic implications (Yoshida et al. 2013, Singh et al. 2015). However, we know little about genome evolution and phenotypic plasticity in asexual plant pathogen lineages and to which degree genetic variation in these lineages affects the virulence profile of pandemic pathogen populations.

*Phytophthora infestans* is a heterothallic oomycete capable of both sexual and asexual reproduction (Judelson and Blanco 2005,Fry 2008). It causes the late blight disease in several plants in the Solanaceae genus *Solanum* and is renowned for its capacity to rapidly overcome resistant host cultivars (Fry 2008,Vleeshouwers et al. 2011). Successive emergence and spread of asexual lineages is a recurrent threat to potato cultivation and one of the main reasons why late blight remains the most destructive disease of potato worldwide (Cooke et al. 2012, Fry et al. 2013, Yoshida et al. 2013,Fry et al. 2015). Interestingly, there is anecdotal evidence that phenotypic variation has emerged within clonal lineages of *P. infestans* with potential consequences on virulence (Forbes et al. 1997,Kamoun et al. 1998).

The asexual lineage EC-1 was first detected in 1990 in Ecuador as a dominant genotype with the A1 mating type and IIa mitochondrial haplotype (Forbes et al. 1997, Yoshida et al. 2013,Yoshida et al. 2014). In Ecuador and neighboring Colombia, EC-1 displaced the pandemic US-1 clone to dominate Andean populations of *P. infestans* associated with cultivated and wild potatoes (Forbes et al. 1997, Oyarzun et al. 1998, Yoshida et al. 2013,Yoshida et al. 2014). As part of a population genomic study of ancient and modern *P. infestans* populations, we previously generated whole genome shotgun sequences of two EC-1 isolates, P13527 and P13626, using Illumina paired-end reads (~51- and 64-fold coverage, respectively) (Tables S1, S2, S3) (Yoshida et al. 2013). Here, we investigated the genetic divergence between the two EC-1 isolates in more detail.

We first estimated levels of presence/absence polymorphisms and copy number variation (CNV) within EC-1. We aligned the sequence reads to the reference genome of *P. infestans* strain T30-4 (Haas et al. 2009) and calculated average breadth of read coverage and read depth per gene as previously described (Cooke et al. 2012). Of the 18,155 *P. infestans* genes, 62 and 60 were missing in the P13527 and P13626 genome sequences compared with T30-4, respectively (Tables S4 and S5). Of these, 12 genes showed presence/absence polymorphisms between the two EC-1 isolates (Table S6). In addition, 69 genes differed in copy number between the two EC-1 isolates (Table S7). Interestingly, P13527 has a much higher number of copies than P13626 for the majority of the genes with CNV (61 out of 69, 88%) (Table S7).

The *P. infestans* genome exhibits an unusual discontinuous distribution of gene density with rapidly evolving disease effector genes localized to expanded, repeat-rich and gene-sparse regions (GSR) of the genome, in contrast to core ortholog genes, which occupy repeat-poor and gene-dense regions (GDR) (Haas et al. 2009,Raffaele et al. 2010). This “two-speed genome” architecture is thought to favor rapid evolution of the pathogen and facilitate adaptation to an ever-changing host environment (Raffaele et al. 2010,Dong et al. 2015). We noted that genes located in GSR were over-represented among the genes with presence/absence polymorphisms and CNV, unlike other gene categories (Figure 1b and d, Figure S1). Genes encoding disease effectors were only over-represented among the genes with presence/absence polymorphisms, unlike other gene categories (Figure 1b, Figure S1). Among the 12 genes with presence/absence polymorphisms within EC-1, 8 are located in GSR, and 7 encode effector genes (Table S6). The missing P13626 genes include 3 genes that have sequence similarity to the RXLR-type *Avr2* effectors, which are detected by immune receptors of the R2 class and, therefore, may reflect adaptation to host immunity (Gilroy et al. 2011,Saunders et al. 2012).

**Figure 1:**
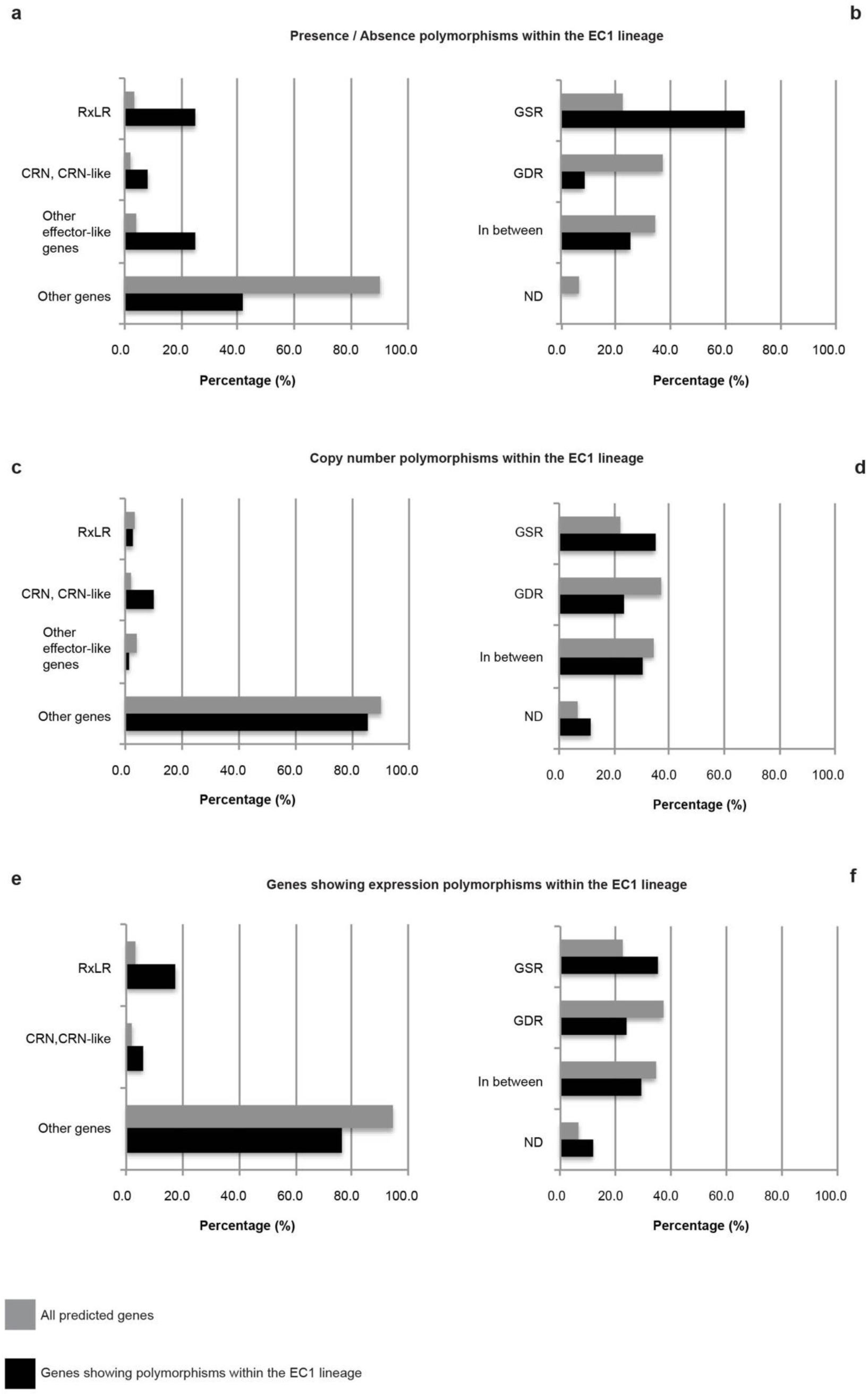
Polymorphisms within the EC1 lineage. **(a)** Percentage of RXLR effector genes, Crinkler (CRN) and CRN-like effector genes, other effector-like genes (including genes coding for small cysteine-rich proteins, protein inhibitors, hydrolases, etc.) and non-effector genes in the set of genes showing presence/absence polymorphisms between P13527 and P13626 compared to that in all the predicted genes. **(b)** Percentage of genes located in Gene Sparse Regions (GSR), Gene Dense Regions (GDR), in between GSR and GDR, and with location not determined (ND) in the set of genes showing presence / absence polymorphisms between P13527 and P13626 compared to that in all the predicted genes. **(c)** Percentage of RXLR effector genes, Crinkler (CRN) and CRN-like effector genes, other effector-like genes (including genes coding for small cysteine-rich proteins, protein inhibitors, hydrolases, etc.) and non-effector genes in the set of genes showing gene copy number (CN) polymorphisms between P13527 and P13626 compared to that in all the predicted genes. **(d)** Percentage of genes located in Gene Sparse Regions (GSR), Gene Dense Regions (GDR), in between GSR and GDR, and with location not determined (ND) in the set of genes showing gene copy number (CN) polymorphisms between P13527 and P13626 compared to that in all the predicted genes. **(e)** Percentage of RXLR effector genes, Crinkler (CRN) and CRN-like effector genes, other effector-like genes (including genes coding for small cysteine-rich proteins, protein inhibitors, hydrolases, etc.) and non-effector genes in the set of genes showing expression polymorphisms between P13527 and P13626 compared to that in all the predicted genes. **(f)** Percentage of genes located in Gene Sparse Regions (GSR), Gene Dense Regions (GDR), in between GSR and GDR, and with location not determined (ND) in the set of genes showing expression polymorphisms between P13527 and P13626 compared to that in all the predicted genes.

Next, we identified SNPs between P13527 and P13626, and compared the SNP frequencies to isolate 06_3928A, which belongs to 13_A2, a distinct clonal lineage of *P. infestans* (Cooke et al. 2012). The number of homozygous SNPs between the two EC-1 isolates was ~54-fold lower than between the EC-1 isolates and 06_3928A, consistent with their expected genetic relationships (Table S8). A large number of SNPs (>327,000) occurred in heterozygous state within each of the individual EC-1 isolates (Table S8). The number of these heterozygous SNPs was ~11-fold higher than homozygous SNPs between the EC-1 isolates and 06_3928A, indicating the maintenance of a high level of genetic diversity at the heterozygous state within the EC-1 clonal lineage (Table S8). About one third (111,144) of the heterozygous SNPs are shared between the EC-1 isolates and 06_3928A, indicating that these sites were already polymorphic prior to the split between the EC1 and 13_A2 clonal lineages (Table S9). The frequency of genes with homozygous and heterozygous SNPs was not dependent on genome localization and functional category (Figure S2).

*Phytophthora* spp. are known to be prone to mitotic loss of heterozygosity (LOH), turning heterozygous genotypes to homozygous in the absence of sexual reproduction (Lamour, Mudge et al. 2012). To estimate LOH, we calculated the number of sites where one of the EC-1 isolates is homozygous whereas the other EC-1 isolate and 06_3928A are heterozygous (Table S9). Assigning 06_3928A as the outgroup established heterozygosity as the ancestral state and LOH within the EC-1 clonal lineage as the most parsimonious explanation. In total, we identified 140,852 heterozygous SNPs that are shared by the two clonal lineages. Of these, 19,983 (14.2%) are homozygous in P13527 and 9,725 (6.9%) homozygous in P13626 indicating significant levels of LOH in EC-1 (Table S9). LOH frequencies did not particularly vary depending on genome localization and functional category (Figure S3). However, we identified 72 effector genes with high or moderate LOH nucleotide substitution effects based on the method of (Cingolani et al. 2012) (Tables S10 and S11). Examples include the predicted RXLR effector PITG_22925 in P13527 and Nep1-like protein PITG_22916 in P13626, which became homozygous for an allele with a premature stop codon, as well as predicted RXLR effector PITG_22922 in P13527 and CRN effector PITG_06050 in P13626, which became homozygous for an allele with multiple nonsynonymous substitutions (Tables S10 and S11).

*P. infestans* is known to exhibit significant levels of gene expression polymorphisms (Cooke et al. 2012). Next, we investigated the degree to which the transcriptomes of the two EC-1 isolates diverge. We isolated RNA from potato leaves 2 days after inoculation with isolates P13527 and P13626, and prepared and sequenced RNA-seq libraries using Illumina technology. We aligned the sequence reads to the *P. infestans* T30-4 transcript sequences and estimated the counts per million (CPM) for each of the 18,155 genes. Gene expression of 13,703 genes was detected in at least one of the two isolates. A comparison of the gene expression between the isolates showed overall similarity (Pearson correlation coefficient = 0.967) (Figure S4).

In total, 17 genes had no detectable RNA-seq reads in one of the two EC-1 isolates but were over the threshold CPM value in the other (see details in Methods). These genes had no fixed nucleotide differences between the two EC-1 isolates in the upstream (<5 kbp), the coding and downstream (<5 kbp) sequences based on alignments to the reference genome (Table S12) (Figure S5). However, of the 17 genes, 12 displayed nucleotide polymorphisms in heterozygous states in one of the two EC-1 isolates (Table S12, Figure S5). Of the 17 genes, 14 genes were expressed in *P. infestans* isolate 06_3928A, indicating that these genes were most likely silenced in the EC-1 lineage (Table S12). Although these overall numbers are small, genes encoding cytoplasmic effectors of the RXLR and CRN families were over-represented among the genes with on/off expression polymorphisms (Figure 1e). However, we did not observe an enrichment in GSR located genes (Figure 1f).

Three among the genes with on/off expression polymorphisms encode RXLR effectors (Table S12). To validate this finding, we isolated total RNA from potato leaves 1 to 5 days after inoculation with P13527 and P13626, and performed RT-PCR analyses (Figure 2). We detected transcripts for PITG_16294 and PITG_23131 in P13527 samples but not in P13626, and transcripts of PITG_01934 in P13626 only (Figure 2). These expression patterns are consistent with the RNA-seq analyses and confirmed that the genes display expression polymorphisms within the EC-1 lineage.

**Figure 2:**
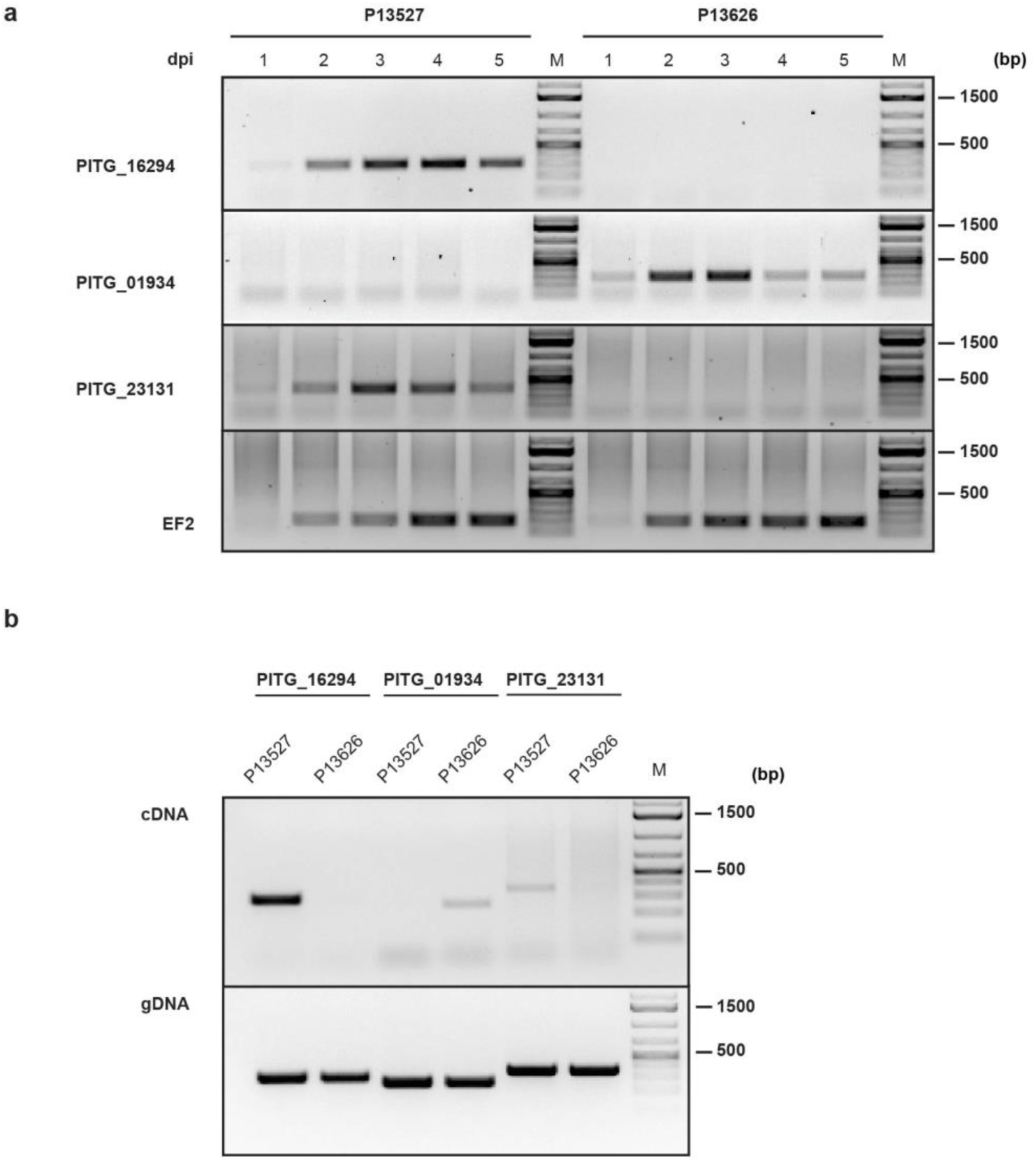
Validation of RXLR effector genes’ expression polymorphisms between P13527 and P13626. **(a)** RT-PCR experiments performed with cDNA samples from infected potato leaves. Samples were collected from 1 to 5 days post-infection (dpi) with P13527 and P13626. Primers specific for the RXLR effector genes PITG_16294, PITG_01934 and PITG_21131 were used for amplification. Amplification of the *EF2* cDNA was included as positive control. ‘M’ indicates the lanes where a DNA molecular weight maker was loaded. **(b)** PCR experiments performed with cDNA (top panel) and gDNA (bottom panel) samples from P13527 and P13626. Primers specific for the RXLR effector genes PITG_16294, PITG_01934 and PITG_21131 were used for amplification. ‘M’ indicates the lanes where a DNA molecular weight maker was loaded.

Plant pathogen effectors with an avirulence activity can trigger hypersensitive cell death when co-expressed *in planta* with their matching immune receptor even in the absence of the pathogen. In an independent set of experiments that aimed at identifying novel *P. infestans* avirulence effectors, we used *Agrobacterium tumefaciens*-mediated transient transformation (agroinfiltration) to express a collection of 240 RXLR effectors in *Solanum venturii* genotype vnt283-1 carrying the *Rpi-vnt1.1* resistance gene (Foster et al. 2009) (Table S13). Six candidates consistently triggered a cell death response typical of hypersensitivity (>70% on a range of 0 (no) -100% (confluent) cell death) (Table S13). Among these, PITG_16294, one of the three RXLR effectors with an expression polymorphism within EC-1, was the only one to trigger cell death only in genotypes that carry the *Rpi-vnt1.1* gene, including in a population of 95 individuals that segregate for this resistance gene (Table S14). To confirm that PITG_16294 activates *Rpi-vnt1.1,* we co-expressed the two genes in the susceptible potato cv. Desiree (Figure 3a). We detected hypersensitive cell death only when PITG_16294 is co-expressed with *Rpi-vnt1.1* or the closely related resistance gene *Rpi-vnt1.2* (Figure 3a). Altogether, these results indicate that PITG_16294 is *Avrvnt1,* the effector that activates *Rpi-vnt1* immunity.

**Figure 3:**
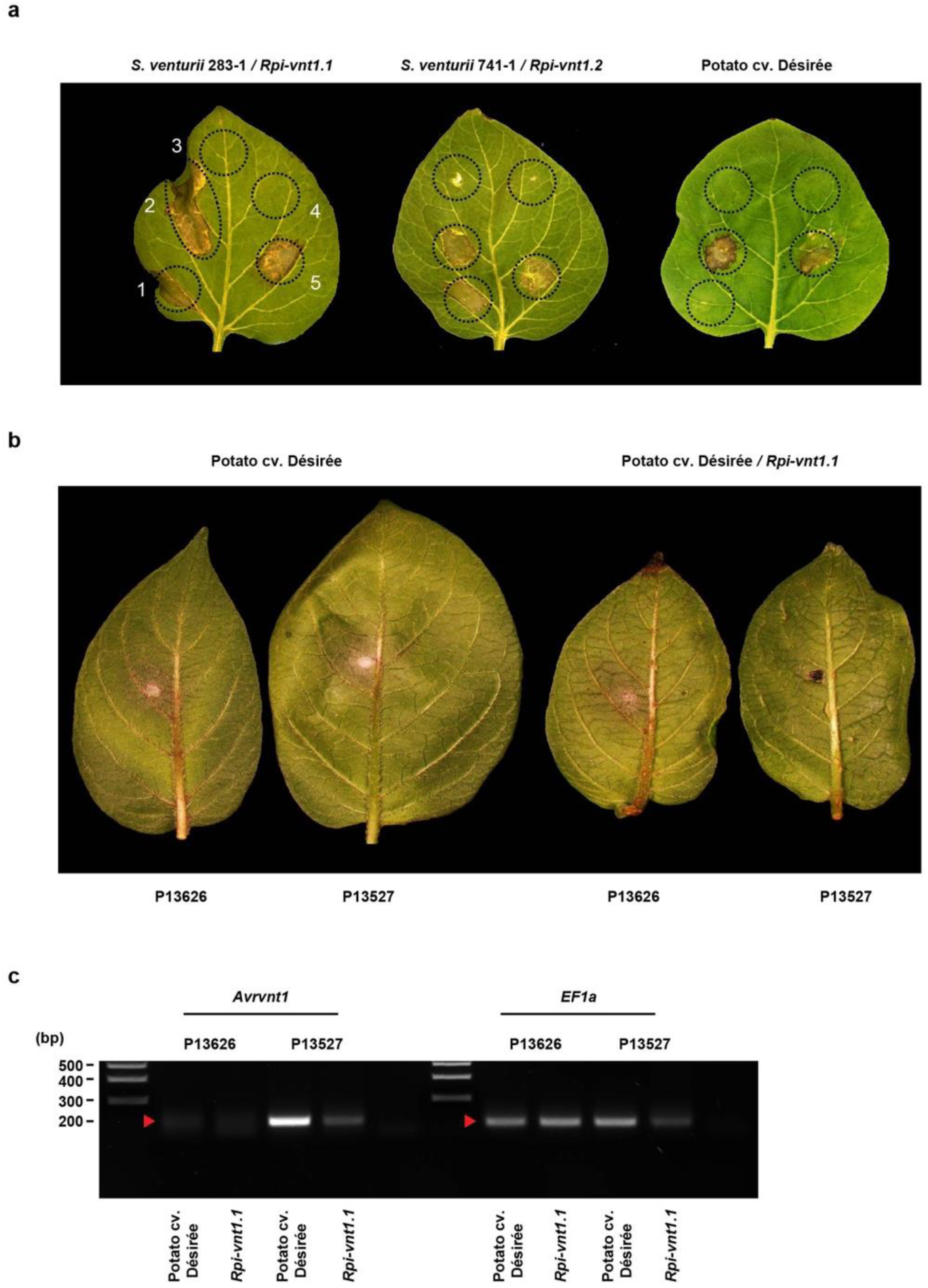
Identification of *Avrvnt1* as the cognate effector for *Rpi-vnt1.* **(a)** Agroinfiltration of *Avrvnt1* leads to cell death in *Solanum venturii* 283-1 and 741-1 carrying *Rpi-vnt1.1* and *Rpi-vnt1.2,* respectively, but not in susceptible potato cultivar Deésireée; (1): *Avrvnt1,* (2): *Avrvnt1* and *Rpi-vnt1.1,* (3): *Rpi-vnt1.1,* (4): empty vector pK7WG2 was used as a negative control in this experiment, (5): *R3a* and *Avr3a* were used as a positive control *R/Avr* pair in this experiment. Note that co-expression of *Avrvnt1* and *Rpi-vnt1.1* in potato cultivar Deésireée (2, leaf on right side) also results in hypersensitive cell death. **(b)** Differential virulence of P13626 and P13527 on plants expressing *Rpi-vnt1.1.* **(c)** RT-PCR experiments performed with cDNA samples from infected potato leaves. Samples were collected at 3 days post-infection (dpi) with P13626 and P13527. Primers specific for the RXLR effector genes PITG_16294, were used for amplification. Amplification of the *EF1a* cDNA was included as positive control.

These results prompted us to test whether the two EC-1 isolates show differential virulence on potato plants expressing the *Rpi-vnt1.1* resistance gene. We used the P13527 and P13626 isolates to inoculate stable transgenic potato cv. Deésireée lines expressing *Rpi-vnt1.1.* The two isolates produced strikingly different phenotypes on these plants (Figure 3b). Whereas P13527 triggered a local immune reaction typical of the hypersensitive response, P13626 infected *Rpi-vnt1.1* potato. Both isolates were virulent on potato cv. Deésireée, which is not known to carry any effective resistance gene against *P. infestans.* RT-PCR on the infected plants confirmed the reduced biomass of the avirulent P13527 on *Rpi-vnt1.1* potato and the absence of PITG_16294 transcripts in P13626 (Figure 3c). We conclude that evasion of *Rpi-vnt1.1*-mediated resistance is probably conferred by absence of PITG_16294 *(Avrvnt1)* transcripts.

Our findings highlight exceptional genetic and phenotypic plasticity in a pandemic asexual lineage of *P. infestans.* We determined that the EC-1 clonal lineage undergoes significant levels of diversification at the sequence and gene expression level. We noted patterns of genome evolution previously described in *P. infestans* even though in this study we examined a shorter evolutionary timescale compared to earlier intra- and inter-specific comparative studies (Haas et al. 2009, Raffaele et al. 2010,Cooke et al. 2012). Therefore, the two-speed genome concept applies to some degree to the relatively recent radiation of the EC-1 asexual lineage, with genes in GSR exhibiting higher levels of structural variation. Linking genome architecture to accelerated adaptive evolution is, therefore, relevant to the microevolutionary timescales observed in agricultural systems. Field pathogen monitoring must, therefore, probe beyond genotyping and genome sequencing to take into account phenotypic plasticity as exemplified by the observed gene expression polymorphisms. In order to fully capture pathogen evolutionary dynamics, approaches based on RNA-seq, such as the recently developed field pathogenomics method (Hubbard et al. 2015,Islam et al. 2016) must be used to complement classical population genetics studies.

We observed that the asexual EC-1 lineage can exhibit phenotypic plasticity in the absence of apparent genetic mutations resulting in a dramatic increase in virulence on a potato carrying the *Rpi-vnt1.1* gene. Such variant alleles may be epialleles that arose through epigenetic changes in the underlying genes as observed in other plant pathogens (Qutob al. 2009, Qutob et al. 2013,Na, et al. 2014). However, it is possible that a genetic trigger, possibly a cis- or trans-structural variant, has resulted in the absence of *Avrvnt1* transcripts. Nonetheless, our observation that on/off expression polymorphisms appears to predominantly affect effector genes points to a potential adaptive process. In the future, understanding the genetic basis of the processes that underlie expression polymorphism and gene silencing is critical to fully appreciate the evolutionary potential of asexual plant pathogen lineages (Gijzen et al. 2014,Soyer et al. 2014).

## Materials and Methods

### Genome sequencing and analyses

Genomic DNA extraction from mycelia of *P. infestans* isolates P13527, P13626 and 06_3928A, library preparation and Illumina next-generation sequencing reactions were performed as described in (Cooke et al. 2012). Sequencing reads with abnormal lengths as well as reads containing Ns were removed with in-house scripts. Filtered Illumina reads were aligned to the reference genome sequence of *P. infestans* T30-4 (Haas et al. 2009) with the Burrrows-Wheeler Transform Alignment (BWA) software. Average breadth of read coverage, read depth per gene and gene copy numbers were calculated as described in (Cooke et al. 2012). Genes with no coverage were considered absent. Copy numbers were estimated from the Average Read Depth (ARD). ARD for the coding sequence of a gene ‘g’ (ARD[g]) was calculated and adjusted according to GC content. Adjusted ARD for a gene ‘g’ (AARD[g]) belonging to the i^th^ GC content percentile was obtained by the formula AARD[g] = ARD[g] * mARD / mARD_GC_ where mARD is the overall mean depth of all coding sequences and mARD_GC_ is the mean depth for genes in the i^th^ GC content percentile. Copy Number for a gene ‘g’ (CN[g]) was calculated as AARD[g] / mARD. Copy Number Variation for a gene ‘g’ (CNV[g]) was calculated as CN[g] in P13527 - CN[g] in P13626. Genes with the same CN in both isolates will have CNV[g] = 0. Genes with CNV[g] ≥ 1 or CNV[g] ≤ -1 where considered as having copy number polymorphisms between P13527 and P13626.

We used the same set of homozygous and heterozygous SNPs in Yoshida et al. (2013). SNPs were called by comparing each isolate with the T30-4 genome sequence. Allele counts for each position were estimated using pileup from SAMtools (Li et al. 2009). SNPs were called as high quality if depth coverage of reads was >= 10. If a high-quality SNP had been detected in the other strains, SNPs with depth coverage of reads >= 3 were rescued. Allele concordance >= 80% was regarded as homozygous SNPs, and allele concordance between 20-80% was regarded as heterozygous SNPs. The details of SNP calling were described in Yoshida, et al. (2013). Synonymous and nonsynonymous (missense and nonsense) SNPs were estimated using snpEff (Cingolani et al. 2012).

To estimate how many homozygous genotypes in segregating sites between P13527 and P13626 were generated by LOH, we calculated the number of segregating sites where one of the EC1 isolates has homozygous genotypes and the other EC1 isolate and the Blue13 isolate 06_3928A have heterozygous genotypes. By using the Blue13 isolate as an outgroup, heterozygous genotypes were estimated to be the ancestral type and homozygous genotypes were predicted to be the novel type generated by LOH in population of EC1 isolates.

### RNA sequencing and analyses

Total RNA was extracted from infected potato leaves at 2 days after inoculation using the Plant RNeasy Kit (QIAGEN). Infection methods followed for this work are described below. Two micrograms of total RNA were used for construction of cDNA libraries using the TruSeq RNA Sample Prep Kit (Illumina) according to the manufacturer’s instruction. The libraries were used for pared-end sequencing with 2 × 76 cycles on Illumina Genome Analyzer IIx (Illumina Inc). Adapter sequences were removed from sequencing reads using cutadapt. Sequencing reads containing Ns were discarded, and then unpaired reads were removed using the perl script “cmpfastq_pe” (http://compbio.brc.iop.kcl.acuk/software/cmpfastq_pe.php).

Paired sequencing reads were aligned to the P13626 consensus transcript sequences using Bowtie2 version 2.1.0. To remove potential PCR artifacts, duplicates of paired sequencing reads, which were exactly mapped to the same positions, were removed using MarkDuplicates of Picard version 1.113. Reads mapped to each transcript sequences were counted with “samtools idxstats” of SAMtools version 0.1.19. Count per million (CPM) for each transcript was calculated.

To search for ON/OFF gene expression polymorphisms between the two EC-1 isolates, threshold value of CPM was defined based on the comparison between RNA-Seq data and microarray data (Cooke et al. 2012) of the 13_A2 isolate 06_3928A from potato leaves 2 days after inoculation. At first, we estimated cumulative distribution of CPM for the transcripts that showed 2-fold induction in planta compared with mycelia in microarray data (Figure S5). Of these transcripts, 95% had over 18 CPM, and then 18 CPM was set as thresholds of expression of genes when the genes showed zero CPM in one of the isolates. This means that when the gene showed zero CPM in one of the isolates and less than 18 CPM in the other, the gene was not regarded as an on/off gene. For the scatter plot, variance stabilizing transformation of the count was calculated using DESeq (Anders and Huber 2010). Subtraction from the transformed value corresponding to CPM=0 from a value of each gene was used as adjust variance stabilizing transformation.

### Enrichment analyses

Gene annotations and locations described in Table S15 were used to categorise genes showing polymorphisms in the genome and transcriptome analyses. Location of genes in Gene Sparse Regions (GSR), Gene Dense Regions (GDR), in between GSR and GDR, and with location not determined (ND) is based on Raffaele et al. (2010). Hypergeometric tests were performed for enrichment analysis to detect effector bias and GSR bias in the sets of polymorphic genes (Supplementary Dataset 1). “phyper” in the R Stats Package was used for the hypergeometric tests.

### Validation of expression polymorphisms

To validate expression polymorphisms in RXLR effector genes, detached Deésireée potato leaves were drop-inoculated with 50.000 zoospores/mL of *P. infestans* P13527 or P13626. Zoospores were harvested from P13527 or P13626 mycelia grown in Rye Sucrose Agar (RSA) plates for 10-12 days at 18°C by adding cold sterile distilled water and incubating for 2 hours at 4°C. The resulting zoospores’ suspension was diluted to 50.000 zoospores/mL with cold sterile distilled water and 10 μL droplets were used to inoculate detached leaves. Inoculated leaves were kept in closed trays with high moisture under controlled environmental conditions (18°C, 16 h of light and 8 h of dark). Leaf disks harbouring the inoculation site were collected at 1, 2, 3, 4 and 5 days after inoculation. For each time point, disks corresponding to different leaflets from five different plants were pooled and RNA was extracted with the Plant RNeasy Kit (QIAGEN). For RNA and genomic DNA extraction from mycelia, P13527 and P13626 were grown in modified plich medium as described in Cooke, Cano et al. (2012). Mycelia grown independently in different plates were pooled and used to perform RNA extraction with the Plant RNeasy Kit (QIAGEN) or genomic DNA extraction with the Omniprep Kit (G-Biosciences). For RT-PCR experiments, 5-10 pg of total RNA from infected leaf tissue and from mycelia were subject to DNAse treatment with the TURBO DNA-free Kit (Ambion). cDNA synthesis was performed with 500 ng of DNAse-treated RNA using the SuperScript III First-Strand Synthesis System (Invitrogen) and oligo-dT primers. cDNA (1 μL) and genomic DNA (200 ng) samples were then used as template in PCRs using DreamTaq DNA Polymerase (Thermo Scientific), following manufacturer’s instructions. Primers used were 5′-TAACGACCCCGACCAAGTTA-3′ and 5′-AGAGATGCCAGCCTTTCGTA-3′ for PITG_16294, 5′-AGTGGTGCTCTCGGCGACTCT-3′ and 5′-AGCCCCTCCGTTTCCTGGGT-3′ for PITG_01934, 5′-ATGCGTCTACCGGTACATGTACGT-3′ and 5′-CTAGTCGTAGTTACGCGTCT-3′ for PITG_23131, and 5′-GTCATTGCCACCTACTTC-3′ and 5′-CATCATCTTGTCCTCAGC-3′ for the *elongation factor-2* control. Presence/absence of amplification products was evaluated in 1 %-1.5% agarose gels and the GeneRuler 1 kb Plus DNA Ladder (Thermo Scientific) was used to assess amplicon size.

### Effectoromics screen and validation of PITG_16294 activity

To identify novel *P. infestans* avirulence effectors a set of 240 RXLR effectors was synthesized at GenScript (New Jersey, USA). Each effector was obtained in pUC57 vector and sub-cloned by Gateway LR cloning into a binary expression vector. The final expression vector was transferred into *Agrobacterium tumefaciens* strain AGL01 and used to perform a screen in *Solanum venturii* genotype vnt283-1 carrying the *Rpi-vntl.1* resistance gene (Foster, Park et al. 2009).

Agroinfiltration of RXLR effectors and/or *R* genes was carried out on wild potato species or Deésireée potato plants according toDu and Vleeshoouwers (Du and Vleeshouwers 2014). The abaxial side of leaves of four-week-old plants was infiltrated using *Agrobacterium tumefaciens* cultures with a final OD_600_ = 0.3. Infiltrated leaves were scored 4 days after agroinfiltration to assess the presence of cell death.

### Assessment of differential virulence on potato plants carrying *Rpi-vnt1*

Fully developed leaves of 10-week old *Rpi-vnt1.1* and wild type Deésireée plants were detached and infected with 10 μL drops of a 50,000 zoospores/mL suspension of *P. infestans* P13527 or P13626, as described above. Inoculated leaves were incubated for 3 days under controlled environmental conditions (18°C, 16 h of light and 8 h of dark). Disks of 1 cm diameter were cut from the areas showing *P. infestans* infection, and used directly for RNA isolation using TRI Reagent (Sigma-Aldrich), according to manufacturer’s protocol. For each sample, 10 μg of total RNA was treated with DNase I (Thermo Fisher Scientific) and 1 μg of treated RNA was used for first-strand cDNA synthesis with a mix of oligo-dT and random hexamer primers, using SuperScript II First-Strand Synthesis System (Invitrogen). Diluted cDNA (1 μL of a 1:5 dilution) was used for PCR with gene specific primers (F: 5′-CGAAGTTGACGGCTCCTG-3′, R: 5′-GGCTCGCTTGAACAAATCC-3′), and with control primers for *elongation factor-1 alpha* (F: 5′-GTCATTGCCACCTACTTC-3′, R: 5′-CATCATCTTGTCCTCAGC-3′). Reactions were performed in 25 μl with 30 amplification cycles (annealing temperature 60°C for both primer pairs) and using homemade Taq polymerase. PCR products were visualized on 1% agarose gel.

## Acknowledgements

This project was funded by the Gatsby Charitable Foundation, Biotechnology and Biological Sciences Research Council (BBSRC), and European Research Council (ERC).

